# Immunological findings of West Caucasian bat virus in an accidental host

**DOI:** 10.1101/2024.09.12.612599

**Authors:** Martina Castellan, Gianpiero Zamperin, Greta Foiani, Maira Zorzan, Maria Francesca Priore, Petra Drzewnioková, Erica Melchiotti, Marta Vascellari, Isabella Monne, Sergio Crovella, Stefania Leopardi, Paola De Benedictis

## Abstract

The *Lyssavirus* genus includes seventeen neurotropic viral species which are able to cause rabies, an acute and almost invariably fatal encephalomyelitis of mammals. Rabies virus (RABV), the genus prototype, is a multi-host pathogen that undergoes multiple events of host-switching, thus occupying several geographical and ecological niches. In contrast, non-RABV lyssaviruses are mainly confined within a single natural host with rare spillover events never followed by adaptation to new accidental host species. In this scenario, unveiling the mechanisms underlying the host immune response against a virus is crucial to understand the dynamics of infection but also to predict the probability of colonization and adaptation to a new target species. Presently, the host response to lyssaviruses has only been partially explored, with the majority of data inferred from RABV infection, under the assumption that all members of the genus exhibit a similar behavior. Through our study we have investigated the immune response determined by the West Caucasian bat virus (WCBV). Indeed, WCBV has been recently associated with a spillover event to a domestic cat, raising concern about the risks for public health due to the circulation of the virus in its natural host. We selected the Syrian hamster as an animal model representative for an accidental host, and chose the intramuscular route in order to mimic the natural route of infection. In hamsters, WCBV was highly pathogenic, determining 100% lethality and mild encephalitis. In comparison with Duvenhage virus (DUVV) and RABV, we found that WCBV displayed an intermediate ability to promote cellular antiviral response, produce pro-inflammatory cytokines, recruit and activate lymphocytes in the hamsters’ central nervous system.

**Author Summary:** Viruses belonging to the genus Lyssavirus cause rabies, a zoonotic and almost invariably fatal encephalomyelitis. However, not all lyssaviruses behave in the same way, with Rabies virus (RABV) being a multi-host pathogen and non-RABV ones remaining mostly confined to their natural hosts. Host-pathogen interaction might help to elucidate the mechanisms behind the differences observed among lyssaviruses in terms of disease dynamics and ability to cross the species barrier, adapting to novel hosts. In our study, we have investigated the host immune response triggered by the WCBV strain spilled-over from its host (the bent-winged bat) to a domestic cat in 2020. Our study offers a key to understand and generalize the mechanisms underlying lyssavirus spillover and pathogenicity in accidental hosts. More generally, our work confirms and marshals previous, fragmentary evidence indicating a strong and inverse relationship between lyssavirus pathogenicity and immune response induction.

## Introduction

Lyssaviruses are the causative agents of rabies, an acute encephalomyelitis of mammals [1,2]. Rabies virus (RABV), the genus prototype, is a multi-host pathogen, with an estimated 59,000 human annual deaths [3–6]. RABV is mainly transmitted through the exposure of non-intact skin or mucosa to the saliva of an infected animal [7,8]. Upon transmission, RABV replicates locally in the wound before spreading from the neuromuscular junction to motor neurons, reaching the peripheral nervous system [8–10] and then the central nervous system (CNS) via microtubule dependent retrograde axonal transport [11–14]. RABV infection is characterized by the inhibition of the host immune response, and indeed, minor inflammatory lesions of the infected brain [15–17].

On the contrary, non-RABV lyssaviruses have restricted geographical and host ranges, mostly associated to bat species, with rare spillover events without the establishment of a new transmission chain [18–21]. Both the pathogenicity and the immunological mechanisms elicited by these viruses have been poorly investigated, with most information inferred from RABV knowledge [22–25], assuming all genus members perform the same. However, increasing evidence highlights the weakness of such an assumption, and indicates that lyssaviruses likely differ in their pathogenicity and stimulation of the host immune response [26–30]. Interestingly, the previous dogma on the invariably lethal infection due to lyssaviruses has also been progressively invalidated [31–33]. As such, i.e. Duvenhage virus (DUVV) has shown little or non-neuroinvasiveness in mice in different experimental setting [34,35] and is able to elicit a strong antiviral response and activation of the interferon (IFN) signalling pathway in mice, similarly to a cell-attenuated RABV strain [36]. Nevertheless, DUVV has been associated to spillover events, with three human deaths reported so far [19,37,38]. Similarly, West Caucasian Bat Virus (WCBV) was first detected in 2002 in a bent-winged bat (*Miniopterus schreibersii*) in Russia [39] and no further identification had been observed until 2020, where a symptomatic WCBV was observed in a cat in Italy [40]. Of note, the reference 2002 WCBV isolate showed to be pathogenic in Syrian hamsters but not in ferrets following intramuscular infection [41]. In this study, we have investigated the pathogen-host interaction and immune response determined by the 2020 WCBV [40] in an accidental host, using the Syrian hamsters as an animal model. To this end, we compared WCBV to RABV and DUVV for which extensive literature is available.

## Results

### WCBV is highly lethal and triggers mild encephalitis in Syrian hamsters

RABV- and WCBV-injected Syrian hamsters all died at 8 and 9 to 15 days post-infection, respectively. The DUVV infection was slightly inefficient, with 5/9 animals (55.55%) developing rabies symptoms (Fig 1A and S1 Fig). We quantified the average level of viral genome in the CNS and observed no significant differences among groups (Fig 1B; S1 Table). All infected hamsters developed encephalitis, characterized by perivascular cuffs of lymphocytes, macrophages, and few plasma cells, with minor inflammatory cell infiltration of the brain parenchyma and leptomeningeal involvement (Fig 1C; S2 Table). Inflammatory changes were observed in the grey and white matter, mainly located in the pons, medulla oblongata, midbrain, thalamus, and the ventral cerebellum. WCBV-infected animals exhibited mild inflammatory changes, with no significant difference in histopathological score compared to RABV, while DUVV exhibited significantly higher scores (Fig 1C and S1 Table). No significant differences were observed the three assessed macro-regions (pons/medulla oblongata, midbrain/thalamus, and cerebellum) for any of the viruses.

**Fig 1.**
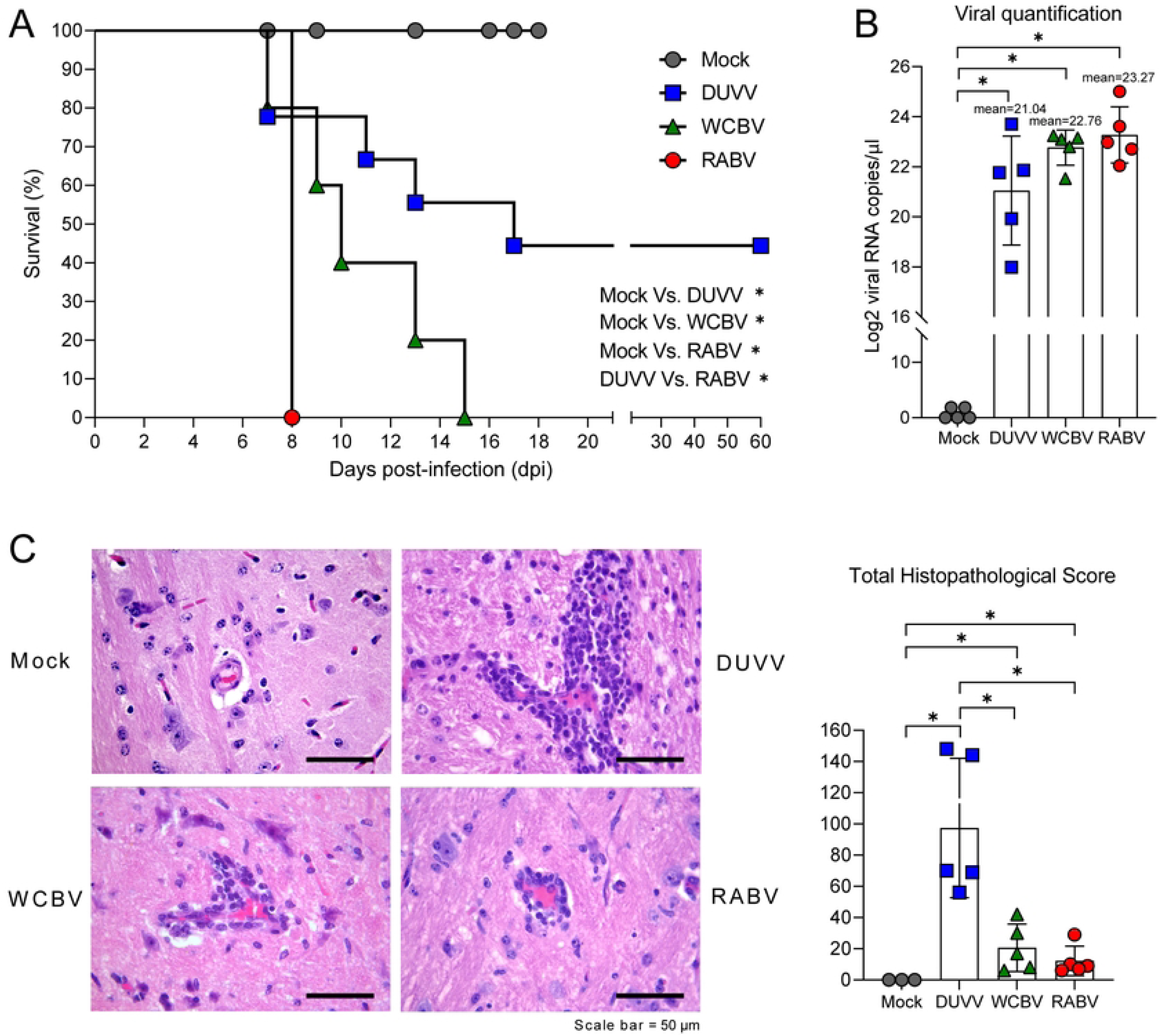
Lyssaviruses infection in the Syrian hamster model and RNA-Seq expression profiles. **(A)** Percent survival over time in Syrian hamsters infected with either DUVV, WCBV or RABV. Kaplan– Meier survival curves are shown by plotting percent survival against time (days post-infection). Mantel–Cox test was performed to compare survival curves of the Mock group *Versus* infected groups and DUVV *Versus* WCBV *Versus* RABV. **(B)** Lyssaviruses viral load as determined through quantitative real time RT-PCR on infected brain samples; results are expressed as Log2 copies/µl of genomic RNA. All the symptomatic animals were analyzed. Mean values ± SD are represented. Wilcoxon–Mann–Whitney test of Mock group *Versus* infected groups and DUVV-*Versus* WCBV-*Versus* RABV-infected hamsters. **(C)** (Left) Representative images of brains from infected and control (Mock animal, pons, animal n. M1, total histopathological score 0) Syrian hamsters; incomplete (mild, score 1) mononuclear perivascular cuffs from the pons of a WCBV-infected animal (N. 6, total histopathological score 30) and RABV (animal n. 2, total histopathological score 9). Severe (score 3) perivascular cuff, composed of more than two cell layers, in a DUVV-infected animal (N. 12, total histopathological score 148). Scale bar = 50 µm. (Right) Final histopathological score of brain lesions of all the symptomatic animals; details on the scores are available in S2 Table. Wilcoxon–Mann– Whitney test of Mock group *Versus* infected groups and DUVV-*Versus* WCBV-*Versus* RABV-infected hamsters. For all the data presented, statistically significant comparisons are shown and represented with a *. P values are listed in S1 Table.

### Lyssaviruses promote a cellular anti-viral state and differently stimulate immune cells recruitment in the CNS

The investigation of the enriched biological processes from the Differentially Expressed Genes (DEGs) through Gene Ontology (GO) analysis (S3 and S4 Tables and S2 Fig) indicated that many processes related to the host response to viral infection and to the activation of the innate immune response were commonly triggered by the three viruses, with DUVV showing the highest enrichment scores (S3 and S4 Tables and Fig 2A). The three lyssaviruses up-regulated genes associated with the cellular anti-viral response (e.g. *Ddx58*, *Mda5* and *Mx1*) and several pro-inflammatory cytokines genes (e.g. *Il1α*, *Il1β*, *Il6*). DUVV also up-regulated genes involved in the NFkB pathway (e.g. *Nfkb1-2* and *Myd88*) and additional pro-inflammatory cytokines-related genes such as *Il12b* or *Il7* (Fig 2B and S3 Table). Real time PCR showed that DUVV-infected hamsters expressed high levels of the cytosolic ss-RNA sensor *Ddx58*, and both DUVV and WCBV induced a greater expression of the pro-inflammatory cytokine *Il1β* (Fig 2C and S3 Table).

**Fig 2.**
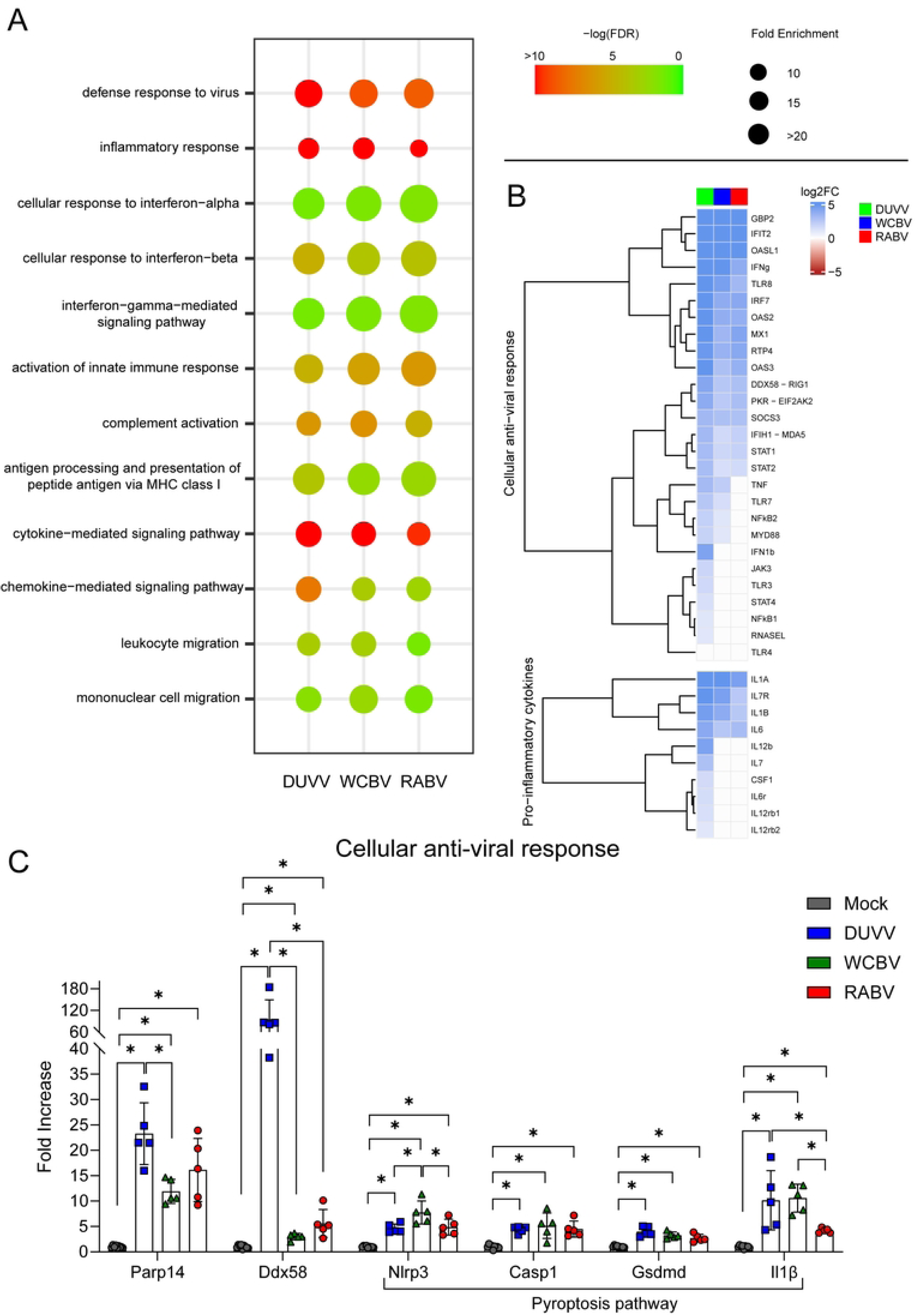
Transcriptomic profile of lyssaviruses-infected Syrian hamsters. **(A)** Dotplot representing the commonly enriched Gene Ontology (GO) terms related to immunity in the lyssaviruses-infected brains. Statistically significant enrichments (FDR < 0.05) are presented and the −LogFDR is shown. See also S4 Table for the raw data on the GO analysis. **(B)** Heatmap (Log2FC values of the infected vs mock comparisons) of selected genes related to the cellular anti-viral response and pro-inflammatory cytokines production. A DEG was significant in a comparison when Log2FC ≤ −1 or Log2FC ≥ 1and FDR < 0.05. See also S3 Table for the raw data on the DEGs analysis. **(C)** Real time PCR results. Relative mRNA expression levels were evaluated as fold increase compared to the mean of the mock animals for each gene analysed. Bars represent mean ± SD and all the single values are also plotted. Wilcoxon–Mann–Whitney test of mock animals *Versus* DUVV-, WCBV- and RABV-infected animals and of DUVV-*Versus* WCBV-*Versus* RABV-infected hamsters. Only statistically significant comparisons are shown and represented with a *. P values are listed in S1 Table.

Moreover, we found that DUVV and WCBV promoted a higher enrichment of “lymphocyte-mediated immunity”, “T cell activation” and “cytokine production” biological processes compared to RABV (Fig 3A). Specifically, DUVV-infected animals fostered “lymphocyte mediated immunity” and “T cell activation” stronger than the other two and exclusively enriched “T and B cell receptor signalling pathway” (S4 Table). We observed that many cytokine-related genes (e.g. *Cd40*, *Cd40l*, *Cd4*, *Cd8b*, *Cd27* and *Cd38*) and genes linked to the microglia activation (e.g. *Aif1*, *Fcer1g*, *Cd40*, *Cd68* and *Cd80*) [15,42] were strongly up-regulated in DUVV- and, to a lesser extent, in WCBV-infected hamsters, with almost zero expression with RABV (Fig 3B and S3 Table). Of note, Syrian hamsters infected with DUVV and WCBV promoted the highest expression of genes related to lymphocyte differentiation (*Cd45*), T- (*Cd3*, *Cd4* and *CD8*) and B-lymphocyte activation (*Cd27* and *Cd38*), with two DUVV-infected hamsters showing the greatest expression levels (Fig 3C).

**Fig 3.**
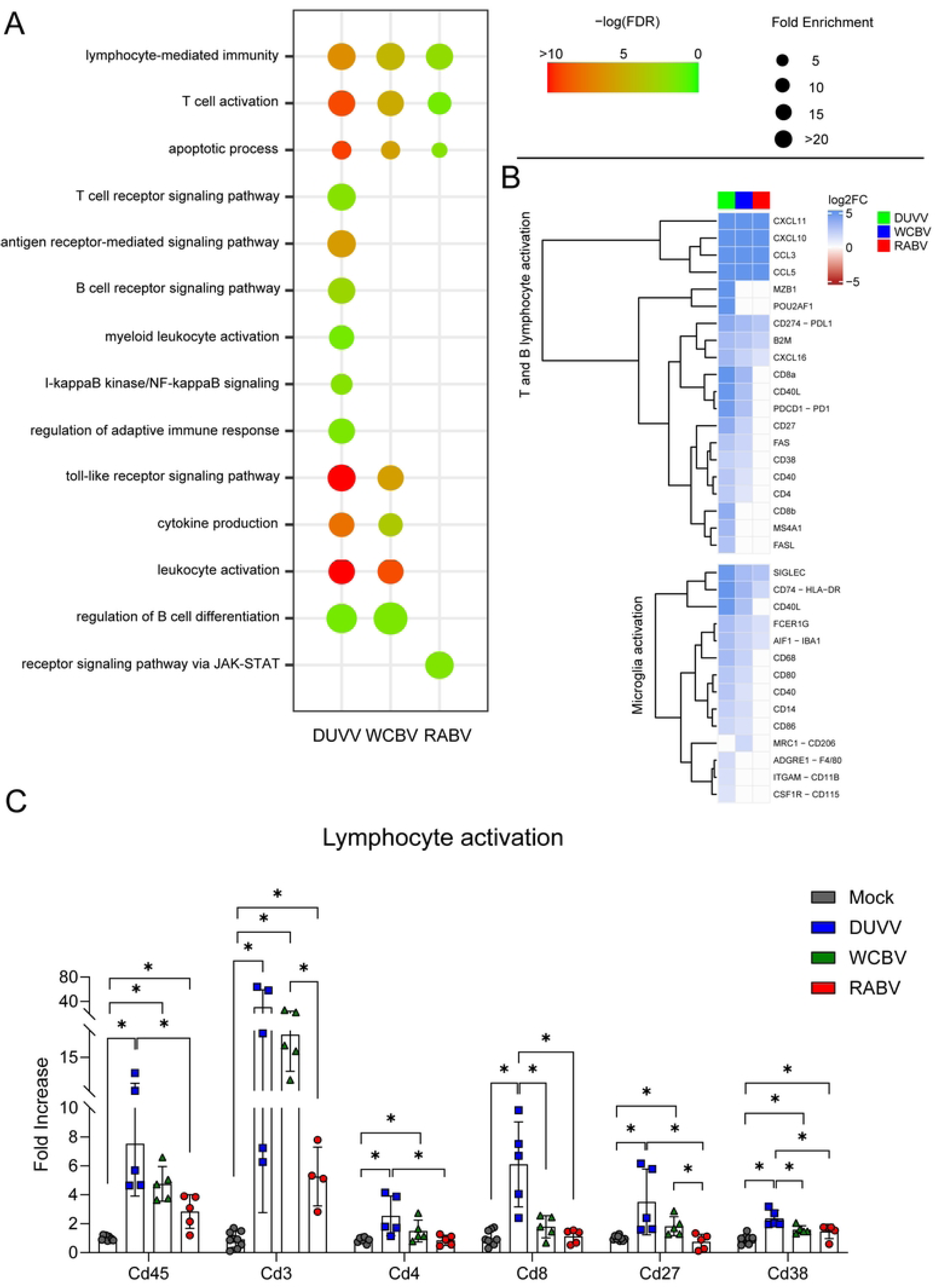
Transcriptomic analyses highlight non-RABV lyssaviruses as being strong activators of the adaptive immune response. **(A)** Dotplot representing the most specific enriched Gene Ontology (GO) terms related to immunity in the lyssaviruses-infected brains. Statistically significant enrichments (FDR < 0.05) are presented and the −LogFDR is shown. See also S4 Table for the raw data on the GO analysis. **(B)** Heatmap (Log2FC values of the infected vs mock comparisons) of selected genes related to the lymphocyte activation and microglia recruitment. A DEG was significant in a comparison when Log2FC ≤ −1 or Log2FC ≥ 1and FDR < 0.05. See also S3 Table for the raw data on the DEGs analysis. **(C)** Real time PCR results. Relative mRNA expression levels were evaluated as fold increase compared to the mean of the mock animals for each gene analysed. Bars represent mean ± SD and all the single values are also plotted. Wilcoxon–Mann–Whitney test of mock animals *Versus* DUVV-, WCBV- and RABV-infected animals and of DUVV-*Versus* WCBV-*Versus* RABV-infected hamsters. Only statistically significant comparisons are shown and represented with a *. P values are listed in S1 Table.

### DUVV stimulates lymphocyte-mediated immunity in the CNS

In all infected brains perivascular cuffs were mainly composed of CD3+ T lymphocytes, Iba1+ macrophages, and a lesser number of PAX5+ B lymphocytes (Figs 4A-C). The number of infiltrating T, B lymphocytes, and microglia/macrophages [43] increased with the severity of histopathological changes. In particular, counts of microglia/macrophages were significantly higher in DUVV-infected animals compared to RABV and WCBV, and counts of T cells were higher in DUVV compared to RABV (Figs 4A and 4B and S1 Table). Interestingly, we noticed that 21 out of the top 30 DEGs were upregulated immunoglobulin fragment-expressing genes in DUVV-infected hamsters (S3 Fig and S3 Table). To confirm what observed as DEGs, we evaluated the presence of IgG immunoglobulins presence [44–46] in the brain of infected and mock animals and confirmed that DUVV had the strongest ability to foster immunoglobulin (Ig) deposition, although the three infections all promoted Ig recruitment in the CNS differently from what observed in mock animals (Fig 4D).

**Fig 4.**
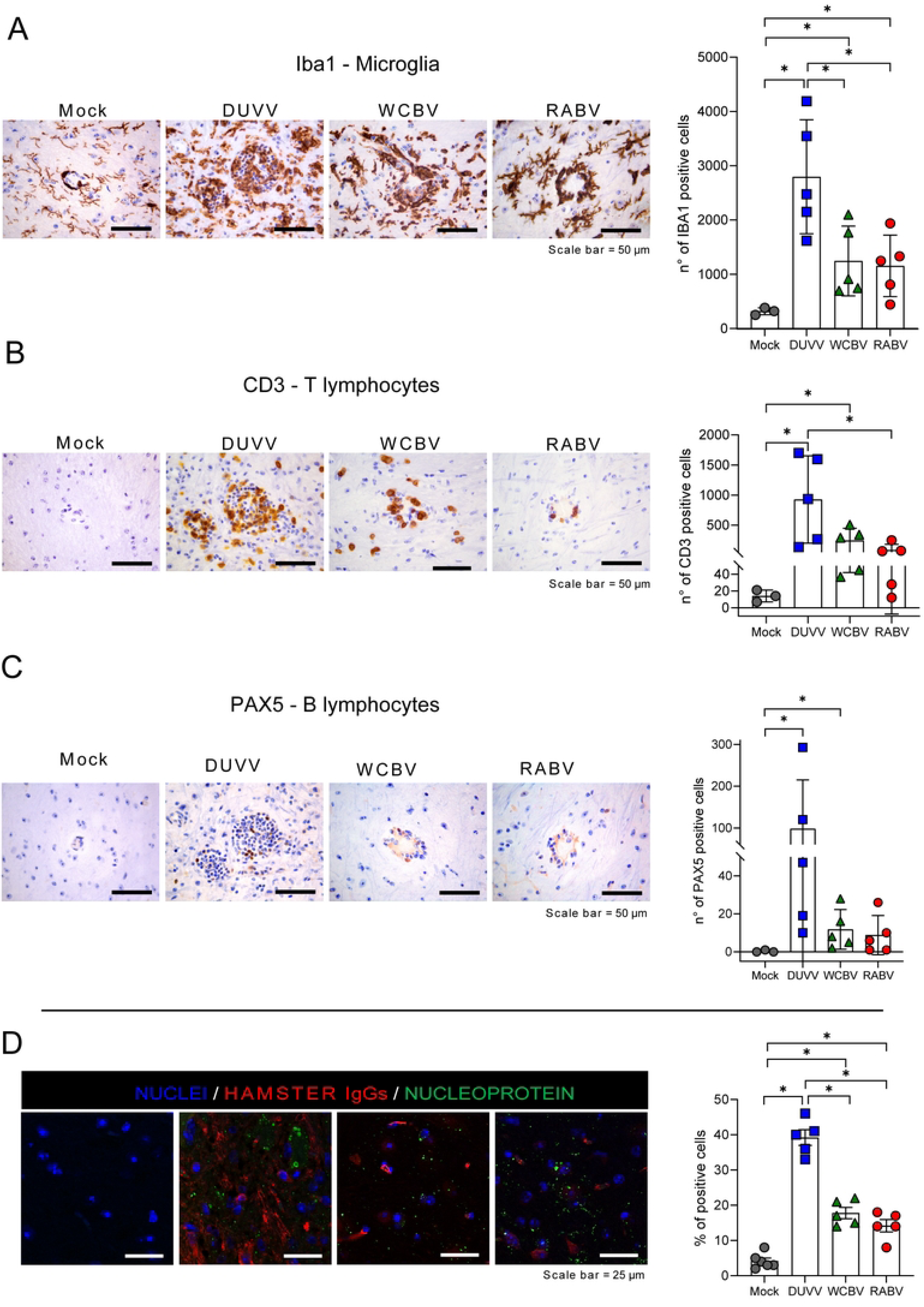
Differential immune recruitment in the lyssaviruses-infected brains. Representative IHC staining (left) for the microglia/macrophage marker Iba1 **(A)**, the T lymphocyte marker CD3 **(B)** and the B lymphocytes marker PAX5 **(C),** representative images of the pons (left). Scale bar = 50 µm. Counts of Iba1-**(A)**, CD3-**(B)** and PAX5-positive cells **(C)** are also shown (right); mean values ± SD are representative of mock group *Versus* infected groups and DUVV-*Versus* WCBV-*Versus* RABV-infected hamsters are shown. **(D)** Immunofluorescence images (left) and relative quantification (right) for hamster immunoglobulin (red) deposition in the infected CNS, coupled with viral Nucleoprotein (green) staining and nuclei (blue). Scale bar = 25 µm. All animals were analysed; representative images are shown. Bars represent mean ± SD of the percentage of positive cells, and all the single values are also plotted. Wilcoxon–Mann–Whitney test of mock animals *Versus* DUVV-, WCBV- and RABV-infected animals and of DUVV-*Versus* WCBV-*Versus* RABV-infected hamsters. Statistically significant comparisons are shown and represented with a *. P values are listed in S1 Table.

## Discussion

The impairment of immune functions is a well-known feature of RABV infection (S4 Fig), which results in fatal outcomes without significant histopathological lesions in the CNS [16,17,29,47]. Until recently, host response against non-RABV lyssaviruses has been improperly inferred from RABV knowledge, despite evidence of lyssaviruses’ differences in pathogenicity. We then investigated the histological changes in the brain and the immune response elicited in the Syrian hamster by a WCBV infection, using the recent 2020 strain.

As expected, our data confirm RABV and DUVV as highly and low pathogenic, respectively [35,36]. Indeed, while RABV caused 100% lethality, DUVV determined the death of only 55% of hamsters (5/9). WCBV was able to invariably reach the hamsters’ CNS where it promoted a productive infection and fatal outcomes in 100% of the infected animals (5/5), mirroring the infection with RABV. This data is consistent with previous preliminary evidences obtained with the 2002 isolate in the same animal model at a similar dose, although a full phenotypic characterization of this WCBV strain was actually missed [41]. WCBV induced mild encephalitic changes, similar to those observed with RABV. Conversely, as expected, DUVV induced severe inflammatory changes, similar to other non-RABV species [34,35,48–50]. Overall, our results support the inverse correlation postulated elsewhere between the lethality of lyssaviruses and the severity of encephalitic lesions [47].

Considering inflammation, RABV and DUVV infections expectedly determined the lowest and the highest ability, respectively, to promote exogenous RNA sensing response, interferon response and pro-inflammatory cytokines production [51,52] (Fig 2A and 2B). Of note, we observed that the cellular antiviral sensing pathway was consistently overexpressed in DUVV-infected animals, confirming DUVV efficiency to stimulate host response (Fig 2B and 2C). The expression levels of genes involved in cellular response to WCBV infection were similar to what observed following the infection with RABV (Fig 2B and 2C), in accordance with the high lethality observed for WCBV and RABV. Of interest, WCBV retained the ability of promoting pyroptosis similarly to DUVV, with a substantial over-expression of both *Nlrp3* and ultimately *Il1β*. Regarding DUVV, our data are in partial agreement with the comparative transcriptomics work by Koraka and colleagues [36], and therefore authors acknowledge that one cannot exclude that the observed findings are related to the strain itself rather than to a viral species peculiarity.

Consistently with the literature, the pro-inflammatory cytokines and chemokines involved in the microglia recruitment and markers of active state [42,53] (Fig 3B) were abundantly up-regulated by DUVV and poorly regulated by RABV. T and B cell recruitment and lymphocyte-mediated immunity was absent in RABV-infected brains while highly promoted by DUVV infection. WCBV fell in between the two extremes, thus retaining a moderate capacity to promote glial activation, and to establish an active pro-inflammatory environment. Based on RNA-seq results and IHC on brain sections, WCBV triggered T lymphocyte and microglia/macrophages recruitment in the CNS, but poorly stimulated B cell activation [54,55] (Figs 3 and 4). Although Ig-related genes’ expression and IgG’s deposition were found in the three infections, only DUVV displayed a strong ability to foster B-cell priming and IgG’s deposition. We speculate that the high stimulation of humoral immunity might explain DUVV low pathogenicity, leading to a more frequent viral clearance.

Altogether, these findings indicate that WCBV seems to behave halfway between DUVV and RABV, promoting both a discreet activation of the anti-viral pathways and the recruitment of the innate and adaptive immunity; these, however, are not efficient weapons to reduce its pathogenicity.

The wide circulation of WCBV in the bent-winged bat (Leopardi et al., under review), the increased opportunity for human encroachment with wildlife [40,56] and the inefficacy of currently available biologicals against this divergent lyssaviruses [57,58] all raise concern about the possibility for this virus to determine with increasing frequency a fatal disease in accidental hosts.

## Material and Methods

### Viruses and Animal experiments

The study involved 28 eight-week-old female Syrian hamsters divided into 5 experimental groups inoculated intramuscularly with either WCBV, DUVV, RABV. Two groups of controls were also prepared Details on the viruses and the experiments are available in S1 File.

### Molecular analyses, histology, immunohistochemistry and immunofluorescence

All the extended procedures are available in S1 File.

### Gene expression analyses by RNA-sequencing

RNA-Seq libraries were prepared from total RNA with the Truseq Stranded mRNA library preparation kit (Illumina), following the manufacturer’s instructions. Sequencing was performed on Illumina NextSeq with NextSeq® 500/550 High Output Kit v2.5 (300 cycles; Illumina) in pair-end [PE] read mode. Details are available in S1 File.

### Statistical analyses

All the extended statistical analyses are available in S1 File.

## Data availability

The RNA-seq data generated in this study have been deposited in the SRA database (https://submit.ncbi.nlm.nih.gov/subs/sra/) under accession number PRJNA1068065. Full genome sequences of DUVV, WCBV and RABV are available under Genbank accession numbers PP869293, MZ501949.1 and OQ787037, respectively. Source data for the RNA-Seq analysis (Figs 2 and 3, S2 and S3 Figs) are provided as S3 and S4 Tables and data for histology, immunohistochemistry and immunofluorescence (Figs 1 and 4 and S1 Fig) are provided as supplementary tables and are available from the corresponding author upon request.

## Ethics

Animal studies were performed in compliance with Directive 2010/63/EU of the European Parliament and of the Council of 22 September 2010 on the protection of animals used for scientific purposes. The procedures were authorized by the Italian Ministry of Health (Decrees 115/2014-PR, 344/2021-PR, 515/2015-PR and 491/2020-PR) before experiments were initiated and endorsed by the IZSVe Ethics Committee. Individual housing exceeded the minimum surface required and was in agreement with the ecology, behaviour, and biology of the species. Seven days prior to infection, the animals were acclimatized in individual cages (BCU-2 Rat Sealed Negative Pressure IVC, Allentown Inc, Allentown, NJ, USA; 19.4cm height x 28.5cm width x 39.3cm depth) for Syrian hamsters. Temperature, humidity, and light–dark cycles were fixed (21 ± 3 °C, 50 ± 10%, and lights off 07:00 AM–07:00 PM, respectively) and monitored throughout the study. All animals had *ad libitum* access to food and water throughout the entire study. The environmental enrichment consisted of gnawing blocks, nesting material, and extra sunflower seeds. We guaranteed monitoring of the animals’ health status at least twice per day and established a humanitarian threshold to avoid unnecessary suffering. Animals were either directly purchased by Charles River Laboratories or purchased and bred in house at the Istituto Zooprofilattico Sperimentale delle Venezie under permission no. 2020/0095.

## Acknowledgments

The authors would like to thank Massimo Boldrin, Franco Mutinelli, Maria Augusta Bozza and the team working at the IZSVe animal facilities for their support in breeding and managing the animals. We want to thank Ronald Mura and Lucas Brandao for the support in the development of the strategy for the RNA-Seq analysis and Annalisa Salviato, Alessia Schivo, Sara Petrin, and Arianna Peruzzo for their suggestions and support with wet lab techniques for transcriptomic analyses. We also thank Giorgia Monetti for providing tissue slices for the immunofluorescence and histological analyses. We acknowledge Marzia Mancin for her fruitful advice on statistical analyses. Finally, we are grateful to Francesca Ellero for the English edits to the manuscript.

## Supporting Information

**S1 Table.** Table representing the statistical results referred to Fig 1, Fig 2C, Fig 3C and Fig 4.

**S2 Table.** a. Histological scoring system for perivascular cuffs; summary of perivascular cuff scores, final histopathological scores, and CD3-, PAX5-, Iba1-positive cells counts. b. Histopathological scores and IHC cells counts; raw data.

**S3 Table.** Numbers of DEGs and Log2FC values found for each comparison made in differential expression analysis for the DUVV-, WCBV- and RABV-infected brains.

**S4 Table.** Numbers of significant enriched GO terms and their scores found for each comparison made in differential expression analysis for the DUVV-, WCBV- and RABV-infected brains. GO terms are presented as blocks of the most specific term (i.e. the ones with the highest level) and their parent terms, based on the child-father relationships characterizing the GO graph.

**S5 Table.** Genetic comparison between the WCBV and DUVV batch produced in new-born mice and the reference sequences. The amino acids count starts from the first methionine of the protein sequence.

**S6 Table.** List of primary and secondary antibodies used for immunofluorescence (IF) and immunohistochemistry (IHC) and details of IHC protocols.

**S7 Table.** List of real-time PCR and quantitative real time RT-PCR primers/probes used in the present work.

**S1 Fig.**
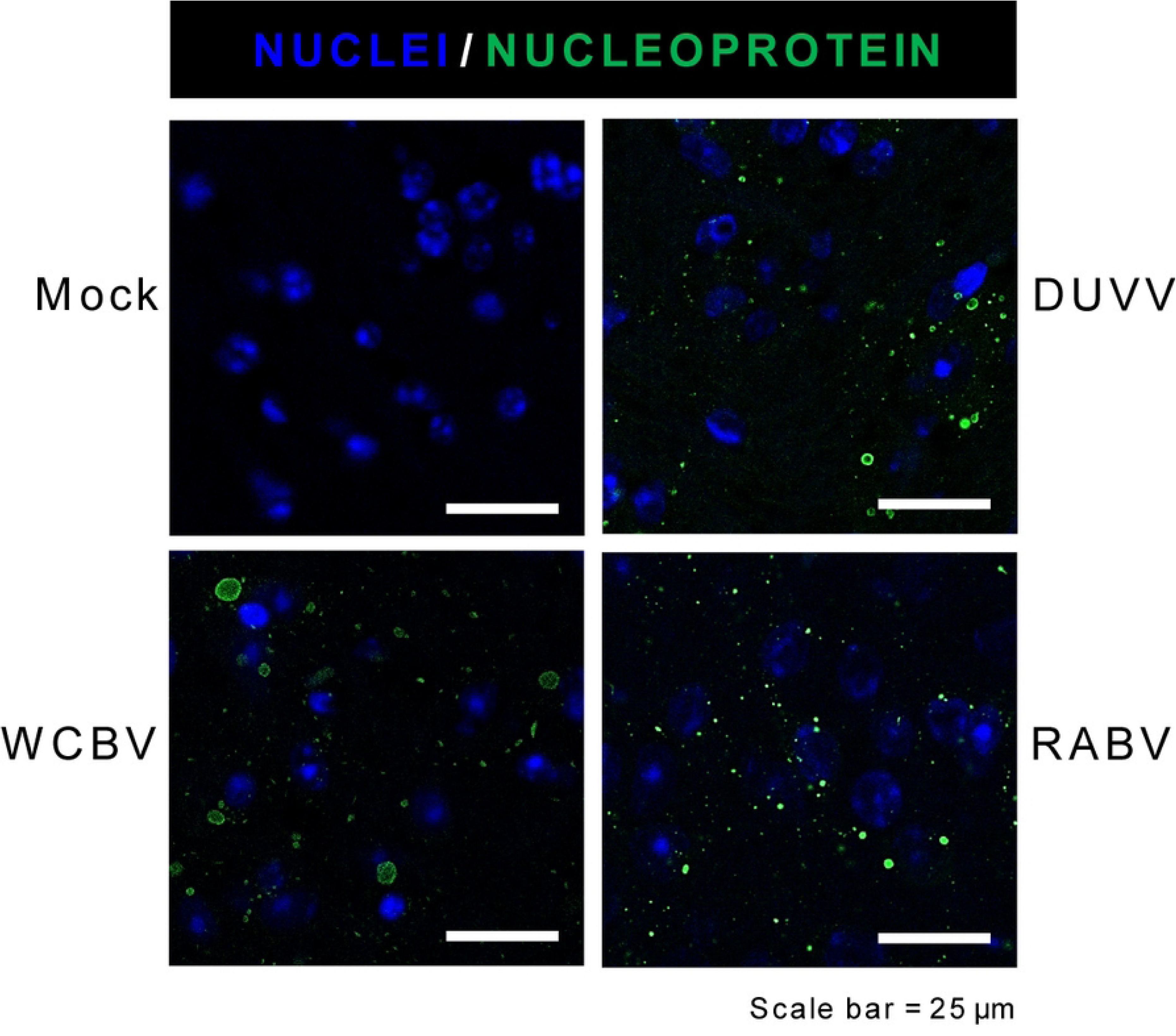
Immunofluorescence staining for lyssaviruses Nucleoprotein (green) and nuclei (blue) in infected hamsters brains. Scale bar = 25 µm. All animals were analyzed; representative images are shown.

**S2 Fig.**
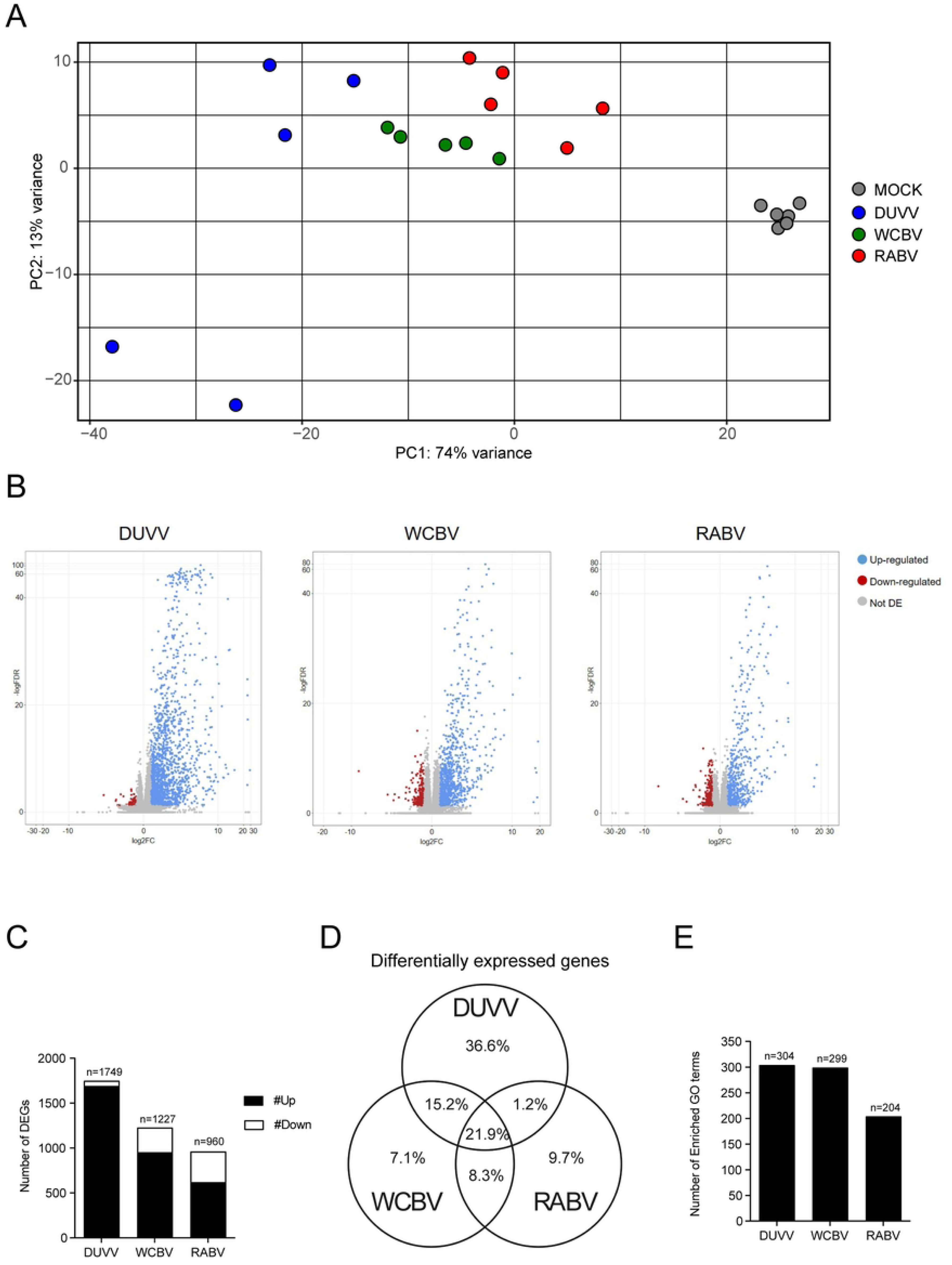
**(A)** Principal component analysis (PCA) of the normalized gene counts of the host genes. The *x* and *y* axes show the two dimensions that explain the overall amount of variance related to gene expression levels. Each replicate is represented by a colored code based on the challenge virus; Mock (grey), DUVV (blue), WCBV (green), RABV (red). The first PCA component (PC1) has the greatest variance (74%), clearly dividing the mock animals from the infected ones, whereas the second PCA component (PC2; 13%) underlines a separation of two samples belonging to the DUVV infection, something that seems to be related to the experiment itself rather than to the specific viral infection. **(B)** Volcano plots showing differential expression analysis results for DUVV-, WCBV- and RABV infected brains (blue, upregulated; red, downregulated; grey, not significant). A DEG was significant in a comparison when Log2FC ≤ −1 or Log2FC ≥ 1and FDR < 0.05. **(C)** Total numbers of DEGs for each comparison of infected *Versus* mock found in the differential expression analysis; up- and down-regulated genes are respectively shown in black and white. DUVV-infected hamsters account for the highest number of DEGs (36.6%) in comparison to WCBC and RABV (7.1% and 9.7%, respectively). See also S3 Table for the raw numbers of DEGs. **(D)** Venn diagram for the graphic representation of the percentages of DEGs associated to each lyssavirus and in common between two or all the viruses. **(E)** Number of enriched GO terms for every comparison of infected *Versus* mock found in the analysis. DUVV and WCBV were characterized by a similar number of enriched GO terms, despite DUVV promoting the different regulation of 40% more genes than WCBV. See also S4 Table for the raw numbers of DEGs.

**S3 Fig.**
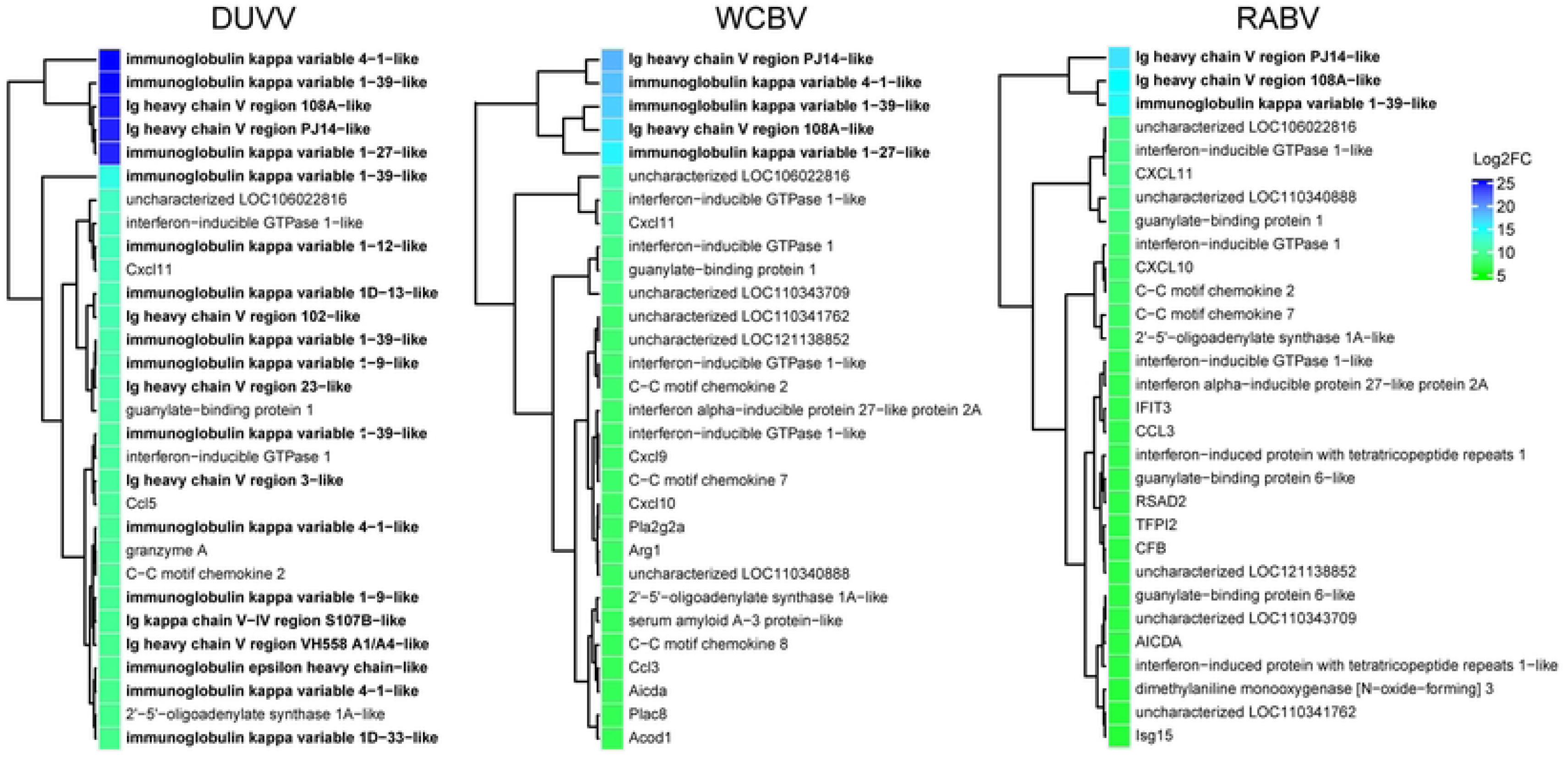
Heatmap (Log2FC values of the infected *Versus* mock comparisons) of the top 30 DEGs in the DUVV-, WCBV- and RABV-infected brains.

**S4 Fig.**
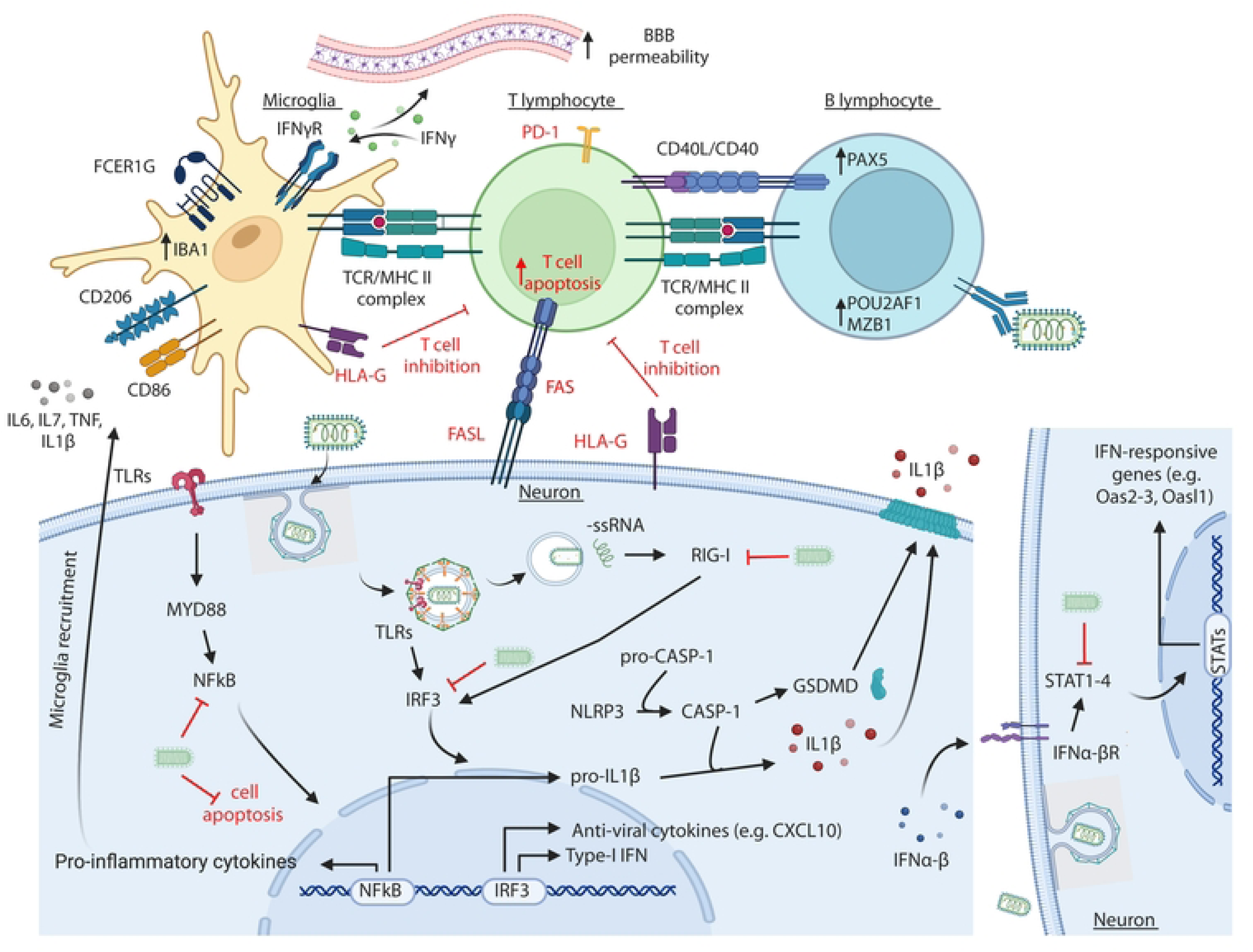
Immune response against RABV in the CNS. Cartoon representing the main cellular pathway and processes that are either activated or inhibited during RABV infection in the CNS. Highlighted in bold and red are the mechanism by which high pathogenic RABV strains blunt the anti-viral and immune response of the host [1,8,16,17,21–23,25,26,28,29,36,42,45,47,53,54,59–88]. Created with Biorender (Agreement number OD26DW39EQ).

**S1 File.** Supplementary Material and Methods.

